# One-stop Microfluidic Assembly of Human Brain Organoids to Model Prenatal Cannabis Exposure

**DOI:** 10.1101/2020.01.15.908483

**Authors:** Zheng Ao, Hongwei Cai, Daniel J Havert, Zhuhao Wu, Zhiyi Gong, John M. Beggs, Ken Mackie, Feng Guo

**Affiliations:** Department of Intelligent Systems Engineering, Indiana University, Bloomington, IN 47405, United States; Department of Physics, Indiana University, Bloomington, IN 47405, United States; Gill Center for Biomolecular Science, and Department of Psychological and Brain Sciences, Indiana University, Bloomington, IN 47405, United States

**Keywords:** Human Brain Organoid, Cannabinoid, Microfluidics, Developmental Biology, Toxicity

## Abstract

Prenatal cannabis exposure (PCE) influences human brain development, but it is challenging to model PCE using animals and current cell culture techniques. Here, we developed a one-stop microfluidic platform to assemble and culture human cerebral organoids from human embryonic stem cells (hESC) to investigate the effect of PCE on early human brain development. By incorporating perfusable culture chambers, air-liquid interface, and one-stop protocol, this microfluidic platform can simplify the fabrication procedure, and produce a large number of organoids (169 organoids per 3.5 cm x 3.5 cm device area) without fusion, as compared with conventional fabrication methods. These one-stop microfluidic assembled cerebral organoids not only recapitulate early human brain structure, biology, and electrophysiology but also have minimal size variation and hypoxia. Under on-chip exposure to the psychoactive cannabinoid, delta-9-tetrahydrocannabinol (THC), cerebral organoids exhibited reduced neuronal maturation, downregulation of cannabinoid receptor type 1 (CB1) receptors, and impaired neurite outgrowth. Moreover, transient on-chip THC treatment also decreased spontaneous firing in microfluidic assembled brain organoids. This one-stop microfluidic technique enables a simple, scalable, and repeatable organoid culture method that can be used not only for human brain organoids, but also for many other human organoids including liver, kidney, retina, and tumor organoids. This technology could be widely used in modeling brain and other organ development, developmental disorders, developmental pharmacology and toxicology, and drug screening.

## INTRODUCTION

Marijuana use has soared in the past decade due to increased public acceptance and relaxation in regulations worldwide. Cannabis use is becoming more popular in pregnancy as a “natural” remedy for symptoms such as morning sickness^1^, with use rate up to 7.47% in pregnant women.^2^ Longitudinal human behavioral studies have shown that prenatal cannabis exposure (PCE) can lead to increased risk-taking and vulnerability to psychosis^3^. However, the exact molecular and developmental impacts of PCE in humans are still largely unknown. Delta-9-tetrahydrocannabinol (THC) is the major psychoactive compound found in marijuana. THC interferes with the endocannabinoid system by acting as partial agonists to CB1 cannabinoid receptors and CB2 cannabinoid receptors. Prenatal THC exposure in animal models has been shown to increase neural progenitor proliferation and decrease neurite growth.^4,5^ Prenatal THC exposure was also shown to compromise astrocyte growth.^6^ Electro-physiologically, prenatal THC exposure decreased spontaneous firing and burst activity of neurons.^7^ These effects combined together could lead to impaired learning and memory in the offspring, as shown in animal models.^8^ Although animal studies have illuminated significant details on the effects of PCE on brain development, the results do not necessarily translate to human subjects since there are profound differences in brain development between lower animals and humans.^9,10^

Recent advances in cerebral organoid have enabled modeling of early human brain development in a dish.^11–13^ Cerebral organoids are human pluripotent stem cell-derived 3D cultures that recapitulate human neocortex development programs. Human brain organoids are excellent models to study human brain development with the following advantages. First of all, human brain organoids intrinsically develop more elaborative subventricular zones (SVZ) with intermediate progenitors as well as outer SVZ (OSVZ) by outer radial glia (oRG). This OSVZ structure is unique to human brain development.^11^ Secondly, the human brain organoid faithfully recapitulates the gene expression and epigenetic programs of prenatal human brains.^14,15^ Human brain organoids can be fabricated from human induced pluripotent stem cells (iPSC), which preserves all genetic makeup and disease-related mutations from human subjects.^16–18^ Lastly, the electrical activity of mature brain organoids can recapitulate human preterm electroencephalogram (EEG) features.^19^ Thus, brain organoids are excellent candidate models to study PCE, to bridge the gap between current animal models and human studies. However, current brain organoid culture protocols are difficult to scale up due to laborious manual manipulation and transferring of organoids during fabrication processes, intrinsic heterogeneity in size, and more importantly, the development of hypoxic cores that prevent their effective use in scalable pharmacology and toxicology studies.

Current brain organoid protocol involves embryonic body (EB) formation in a low adherence or hanging drop plate. The formed EBs are then embedded in the extracellular matrix (ECM) on parafilm or other non-adherent surfaces, followed by growth in a rotation vessel or bioreactor culture. These conventional processes involve transferring brain organoids to at least 3 different vessels and laborious medium change steps during culture processes, which could lead to loss of EB, potential physical damage and even contamination risks. Moreover, since multiple brain organoids were cultured in the same well in a multi-well plate format, brain organoids can merge with each other during the formation process, causing inconsistency in organoid size and structure. Meanwhile, brain organoids can reach over 4 mm in diameter, causing necrosis as well as hypoxic cores, altering gene expression profile during neurodevelopment. To reduce hypoxic core formation, air-liquid interface culturing has been adopted for brain organoid formation to successfully minimize hypoxia and enhance cell survival in prolonged cultures.^20^ Moreover, 3D printed bioreactors have been used to improve brain organoid uniformity and throughput, which allow for a simultaneous culture of 32 organoids each time.^16^ However, there are still tremendous unmet needs to develop new technologies for high throughput generation of brain organoids without huge size variation, necrosis, and hypoxic cores.

Organoids-on-a-chip technologies provide advantages to recapitulate organ physical structures, chemical environment, gas exchange as well as mechanical cues^21–23^. Ingber group has developed a lung-on-a-chip device consisting of two cell culture channels and an interface membrane, and this device design can precisely control fluidic flow and mechanical deformation to mimic lung breathing as well as intestinal epithelium stretching.^24–29^ Moreover, the microstructure-based perfusion chip design also allows for robust generation of organoids and precise control of medium perfusion, and this device has been adopted to fabricate liver, pancreas and glomerulus organoids and even full embryoids on chip.^30–43^ Body-on-a-chip designs feature interconnected modular organoids to mimic inter-organ crosstalk.^44–48^ To address the challenges in human brain organoid culture, the Qin group recently has developed a series of microfluidic chips to simplify the brain organoid fabrication process and minimize transferring loss of brain organoids. Micropillar array devices have been used for the *in situ* generations of human brain organoids.^49^ A perfusable organ-on-a-chip system has been developed to culture embryonic bodies which can be further induced into brain organoids.^50^ Moreover, the same group also fabricated hollow fiber to physically restrict organoid size and prevent organoids from merging with each other.^51^ Reiner group has developed a brain organoid chip to investigate the impact of physical force on brain organoid development.^52^ Importantly, increasing attention has been attracted to further develop microfluidic and engineering technologies to simplify the culture procedure, improve the uniformity and reproducibility, provide better geometrical confinement and environmental control of brain organoids.

To address the technological barriers in current brain organoid culture, we developed a novel microfluidic platform with several unique features. (1) This simple device establishes an integrated workflow that enabled a one-stop assembly and culture platform for brain organoids, including *in situ* EB formation, neuroectoderm induction, extracellular matrix embedding, and brain organoid maturation. (2) This device also enhances brain organoid uniformity by physical constriction to avoid the random brain organoid merging and control organoid size. (3) This device incorporates air-liquid interface culture *in situ*, which in combination with the physical size restriction, minimized hypoxic core formation. (4) This device is compatible with current commercially available 6-well or 24-well plate, which can scale up brain organoid fabrication for future studies of human brain development under a wide range of chemical exposure conditions. Using this technology, we investigated the alteration of human neuron development under prolonged THC exposure (**Fig. 1a**). By using this device, we were able to fabricate 169 human brain organoids each batch in a standard 6-well plate format (**Fig. 1b**), and these organoids structurally recapitulated neural development. With an on-chip THC treatment, these organoids reduced neuron maturation, reduced CB1 expression, as well as reduced neurite outgrowth, and the effect of transient THC exposure on brain organoid electrical activity was studied. This is one of the first studies to interrogate the impact of PCE on human brain development in human brain organoid models.

**Figure 1.**
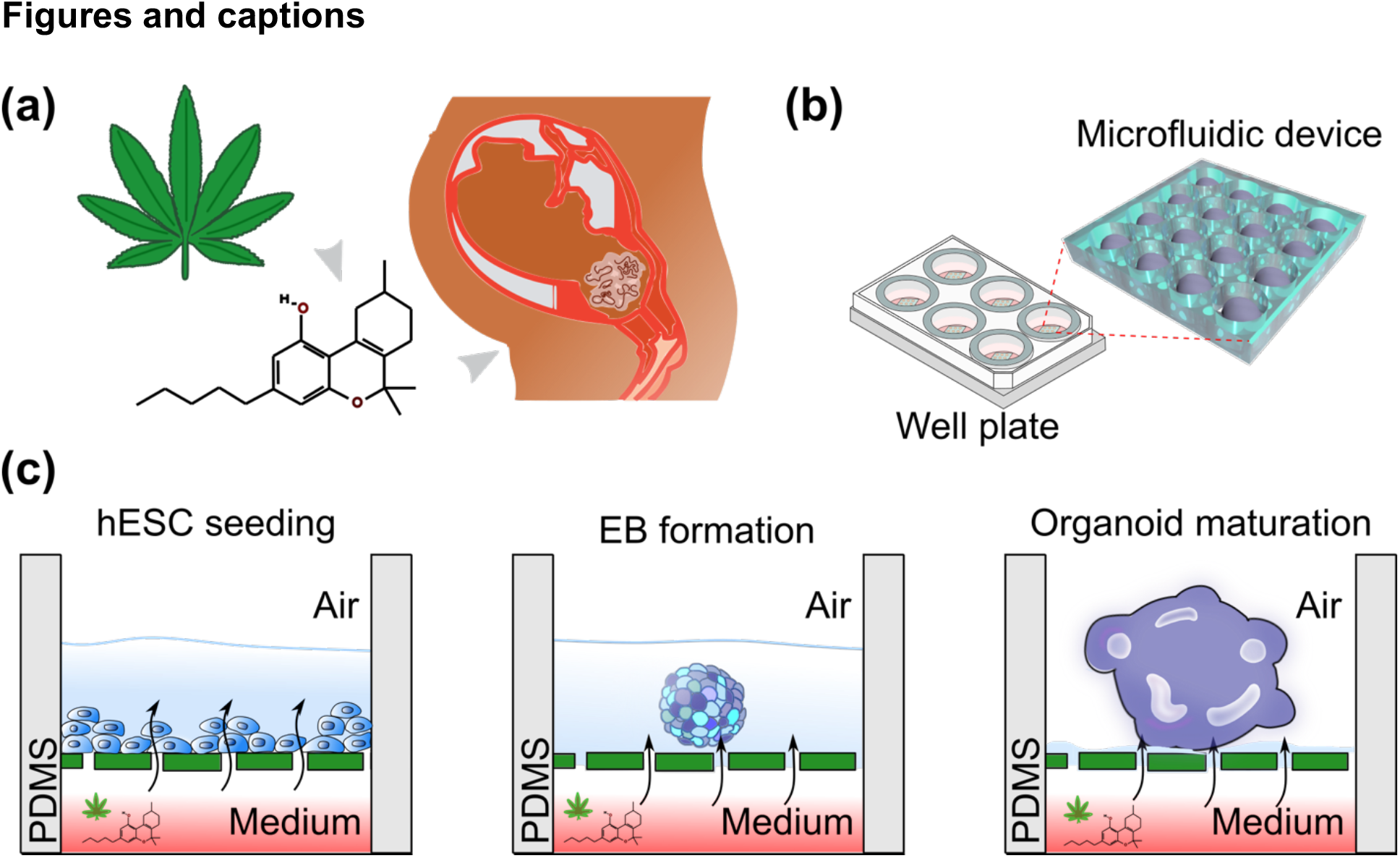
(a) Prenatal cannabis (e.g. cannabinoids such as THC) exposure may impact on fetal brain development. (b) Schematics of a microfluidic brain organoid assembly device that compacts with well plates. (c) Schematics of brain organoid fabrication and cannabis exposure within the microfluidic device.

## EXPERIMENTAL SECTION

### Microfluidic device fabrication

The microfluidic device was fabricated by bonding a polydimethylsiloxane (PDMS) chamber to a hydrophobic polytetrafluoroethylene (PTFE) coated stainless steel metal mesh. Briefly, the PDMS precursor was mixed at 10:1 (w/w) with the curing agent. PDMS was molded into a 3.5 cm x 3.5 cm cylinder block with 5 mm in height, and gas bubbles were removed using a vacuum chamber. The PDMS prepolymer was then cured at 80°C oven for 1 hour. 2 mm diameter perfusable chamber arrays to hold brain organoids were created using a 2 mm needle puncher. A 13 x 13 array of cylinder wells were punched onto each PDMS block. The PDMS blocks with perfusable chambers were then bonded to PTFE coated stainless steel mesh (TWP Inc) using PDMS pre-polymer and thermally cured in an 80°C oven for 1 hour. The whole device was then autoclaved at 121°C for 30 minutes. The autoclaved sterile device was then assembled with a 6 well polyester transwell insert (VWR).

### Human embryonic stem cell culture

Human embryonic stem cell line WA01 was purchased from WiCell. WA01 cells were maintained on Matrigel (Corning) coated 6-well plates, supplemented with mTeSR1 medium (Stemcell Technologies) in a 37°C, 5% CO_2_ supplemented incubator. WA01 cells were passaged using Versene solution (Stemcell Technologies) every 6-8 days.

### Microfluidic assembly of cerebral organoids

Cerebral organoids were fabricated using a stemdiff cerebral organoid kit (Stemcell Technologies). Initially, 9,000 WA01 cells were injected into each 2mm perfusable chamber punched on our PDMS device. With a 6 well transwell insert (VWR), we initially loaded the bottom chamber with 2 milliliters of embryonic body formation medium and layered 1 milliliter of the same medium on top of the PDMS mold. WA01 cells were allowed to spontaneously aggregate and form EB for 4 days. Formed EBs were then subject to neural induction medium for 2 days. On day 7, we loaded each 2 mm perfusable chamber with 20 microliters of Matrigel. Matrigel gel was allowed to solidify in perfusable chambers for 30 minutes. The PDMS mold holding the brain organoid was then topped with a neural expansion medium. At day 10, the toplevel medium was removed, and the bottom chamber medium was reduced to 1 milliliter to create air-liquid interface culture conditions.

### Live/Dead staining

To visualize brain organoid viability, EB and mature organoids were stained using a live/dead cell imaging kit (Invitrogen). Briefly, brain organoids were submerged in medium supplemented with 2 μM of carboxyfluorescein succinimidyl ester (CFSE) and 4 μM of ethidium homodimer (EthD) for 4 hours. The staining medium was then removed and replaced with fresh medium. Organoids were then visualized using an inverted fluorescence microscope (Olympus, IX-81).

### Hypoxia staining

To identify hypoxic core formation, brain organoids were submerged in culture medium supplemented with Image-IT™ hypoxia reagent (Invitrogen) overnight. After replacing the staining medium, brain organoids were left to sit in a fresh medium for an additional 4 hours. The stained organoids were then imaged under an inverted fluorescence microscope (Olympus, IX-81).

### Microelectrode arrays (MEA) measurement

Mature brain organoids at 1 to 3-months of age were incubated on polyethyleneimine (PEI) pre-treated MEA plates with 10 μg/mL laminin (Sigma) for 4 hours to ensure attachment to the plate. Once the brain organoids had attached, we then topped the organoids with 1 mL of fresh BrainPhys neuronal medium (Stemcell Technologies) and incubated them in a 37 °C, 5% CO_2_ supplemented incubator. The organoids were allowed to adhere to the MEA plate for at least 4-6 days before measurements were taken using a multichannel MEA system. Signals were recorded at a sampling frequency of 20 kHz and processed for spike detection using a butterworth bandstop 300-3000 Hz filter. Threshold was set as 4 standard deviation from mean. Subthreshold activity was sorted with activity duration from 0.3-3.0 milliseconds. For drug treatment, delta-9-THC was dissolved in DMSO and dosed at a concentration of 1 μM.

### Cryo-section and immunofluorescence staining

For immunofluorescence staining, brain organoids were washed twice with phosphate-buffered saline (PBS) and fixed in 4%paraformaldehyde at 4°C overnight. Fixed organoids were then transferred to 30% sucrose to cryoprotect overnight at 4°C. Cryoprotected organoids were then embedded in cryomolds (Sakura Finetek) with O.C.T compound (Fisher Healthcare) on dry ice. Embedded brain organoids were sectioned on a cryostat to 20 μm thick slices. Organoid slices were blocked with 5% normal goat serum (Abcam) and 0.3 % Triton™ X-100 (Sigma) in 1x PBS for 1 hour at room temperature. The slices were then incubated with the indicated primary antibodies at 4°C overnight. Respectively, slices were stained with anti-PAX6 (1:500, BioLegend, Catalog 901301), anti-MAP2 (1:500, Millipore, Catalog AB5543), anti-GFAP (1:500, DAKO, Catalog Z0334) or anti-CB1 (1:500, generated in the house as previously reported^53^). After primary antibody incubation, the slices were washed 3 times with 1x PBS. The corresponding secondary antibodies (Invitrogen, 1:500) were then introduced to the slices and incubated at room temperature for 1 hour, followed by 3 times PBS washes and counterstaining of cell nucleus using 4’,6-diamidino-2-phenylindole (DAPI).

### Gene expression analysis

To analyze gene expression in brain organoids, 30-day old organoids were lysed for total RNA extraction using an RNeasy mini kit (Qiagen). Total RNA was then reverse-transcribed using a qScript cDNA Synthesis Kit (Quantabio). We then performed qPCR using SYBR green (Thermo Fisher) with the following primers as previously reported^50^: Triplicate experiments with 3 brain organoids from each treatment group were performed, each qPCR reaction was also done in triplicates. PAX6 forward 5’-AGT TCT TCG CAA CCT GGC TA-3’, PAX6 reverse 5’-ATT CTC TCC CCC TCC TTC CT-3’, SOX2 forward 5’-GGA TAA GTA CAC GCT GCC CG-3’, SOX2 reverse 5’-ATG TGC GCG TAA CTG TCC AT-3’, CTIP2 forward 5’-CAG AGC AGC AAG CTC ACG-3’, CTIP2 reverse 5’-GGT GCT GTA GAC GCT GAA GG-3’, TUJ1 forward 5’-CTC AGG GGC CTT TGG ACA TC-3’, TUJ1 reverse 5’-CAG GCA GTC GCA GTT TTC AC-3’, CB1 forward 5’-CAG TGA AGA GCC TGG GAA GG-3’, CB1 reverse 5’-GGT CAG CAA GTC AGT CCG TC-3’.

### Neurite outgrowth analysis

Mature brain organoids cultured for 30 days were allowed to attach to fibronectin glass coverslips for 4 hours in the presence of 10 μg/mL of laminin (Sigma). Attached brain organoids were then incubated on coverslips for 3 days under DMSO vehicle or THC (1 μM) treatment and stained with celltracker green (Invitrogen) to visualize neurons. Neurite outgrowth and coverage area were imaged under an inverted microscope (Olympus) and quantified using ImageJ software.

### Statistical analysis

Statistical analysis was performed using Student’s T-test. P-values were expressed as *p < 0.05, **p < 0.01, ***p < 0.005.

## RESULTS

### One-stop microfluidic organoid assembly

Brain organoid fabrication involves 4 key steps: embryonic body (EB) formation, neuroectoderm induction, neuroepithelium expansion (extracellular matrix embedding step) and brain organoid maturation. Traditionally, organoids are generally transferred from the EB formation plate to Matrigel embedding parafilm followed by a spinning flask or rotation vessel culture.^54^ To simplify this process and perform all these 4 steps in one single device, we fabricated a PDMS microfluidic device for brain organoid formation, and this 3.5 cm x 3.5 cm circular device consists of 169 cylinder-shaped perfusable chambers (a diameter of 2 mm and a height of 5 mm) and PTFE coated wire mesh as the bottom (**Fig. 1b**). PTFE wire mesh is hydrophobic which prevents hESC adherence and allows suspended hESC to form embryonic bodies spontaneously. Initially, 9,000 dissociated WA01 human embryonic stem cells (hESC) were seeded into each well within a 6 well-plate. Suspended WA01 cells will spontaneously aggregate into embryonic bodies. After the EB formed, we then sequentially subjected it to neural induction and expansion medium, followed by Matrigel embedding in the same microwell, with the brain organoids submerged in medium. Brain organoids were allowed to mature until day 10. We then lowered the culture medium level to an air-liquid interface mode (**Fig. 1c**). This allowed the brain organoid to mature in atmospheric oxygen levels. The brain organoids were maintained in an air-liquid interface culture for 1-3 months before we characterized them.

### On-chip organoid culture

Our microfluidic device improves brain organoid fabrication processes in three aspects: (1) In contrast to the traditional method requiring transfer of brain organoids between hanging drop plate for EB formation, non-adherent plate for suspension culture and parafilm for Matrigel embedding, our method requires only one culture vessel, preventing loss or damage to EB during the transfer process, and avoiding potential contamination risk. (2) Our air-liquid interface minimizes hypoxic core formation. (3) Our microfluidic device physically restricts brain organoid growth and controls their size to be under 2 mm, reducing size variation in brain organoid culture and preventing non-uniform necrosis formation. We imaged these brain organoids at different stages. Brain organoids were stained using a live/dead staining kit, with live cells were labeled with carboxyfluorescein succinimidyl ester (CFSE) green fluorescence and dead cells were labeled with ethidium homodimer-1 (EthD-1) red fluorescence. We could visually confirm that brain organoids fabricated by our devices continued to be highly viable and developed neuroectoderm and matured similarly as compared with the conventional method (**Fig 2**). Moreover, brain organoids fabricated using conventional method reached a diameter >3mm at day 30, creating a significant necrosis core as indicated by ethidium homodimer staining of dead cells. Whereas for organoids cultured on our device, the size was physically restricted to under 2mm. And they were cultured on an air-liquid interface, thus the necrotic cores were minimized (**Fig 2**). To test whether this high viability was a benefit from lowered hypoxic levels in the center of brain organoids, we also stained these organoids with a fluorescent hypoxia indicator. We found that indeed brain organoids’ hypoxic core formation was drastically reduced by microfluidic air-liquid interface culture (**Fig. 3a-b**). In addition, physical restriction wells could minimize organoids merging and size heterogeneity. As shown in **Figure. 3c-d**, brain organoid size became more uniform around 2 mm as compared with control cultures, where many merged organoids, reaching 4 mm in the longest diameter were generated during rotating vessel culture.

**Figure 2.**
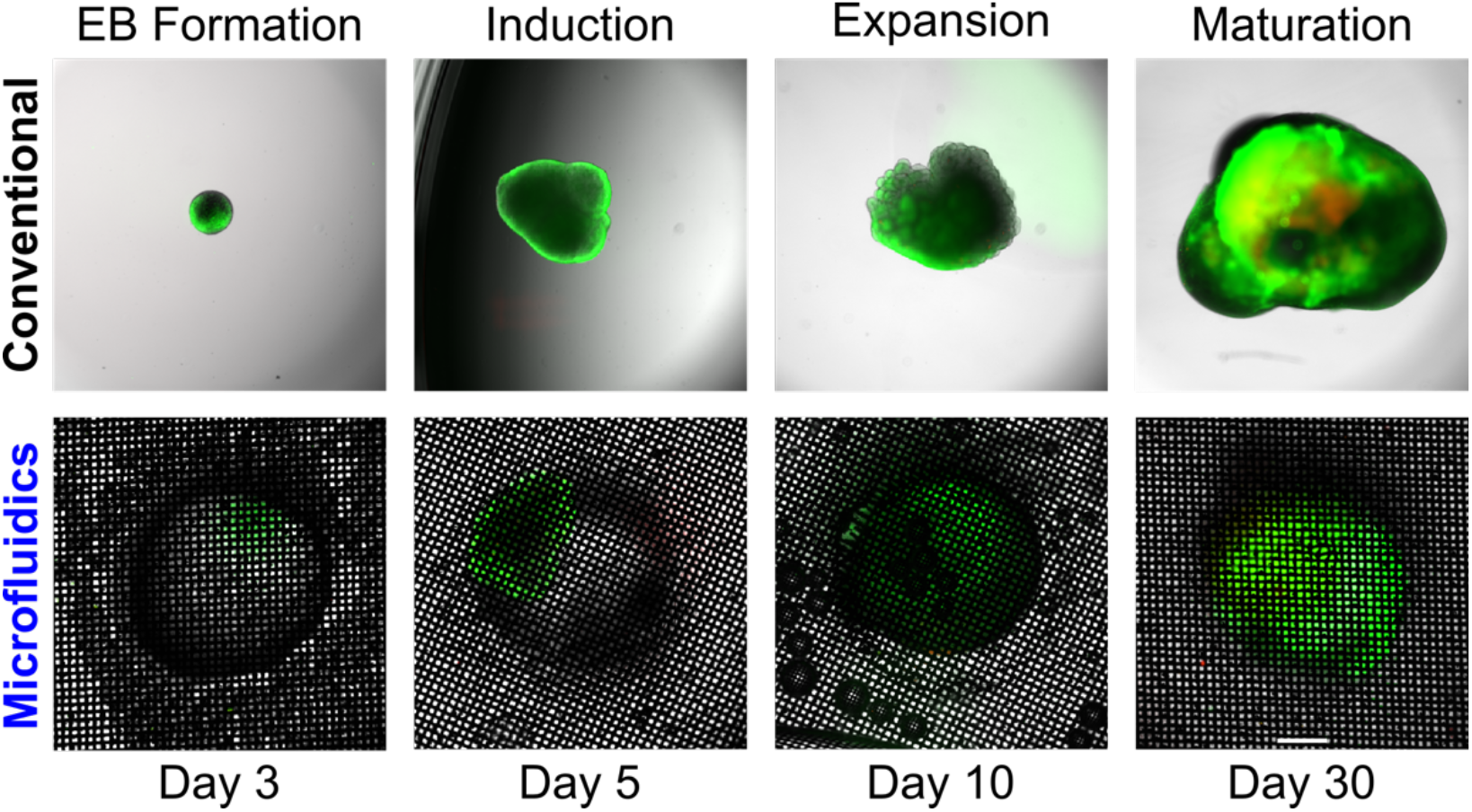
Comparison of cerebral organoid fabrication process using the conventional protocol and microfluidic method. Organoid viability was visualized by live(green)/dead(red) staining. Scale bar: 1 mm

**Figure 3.**
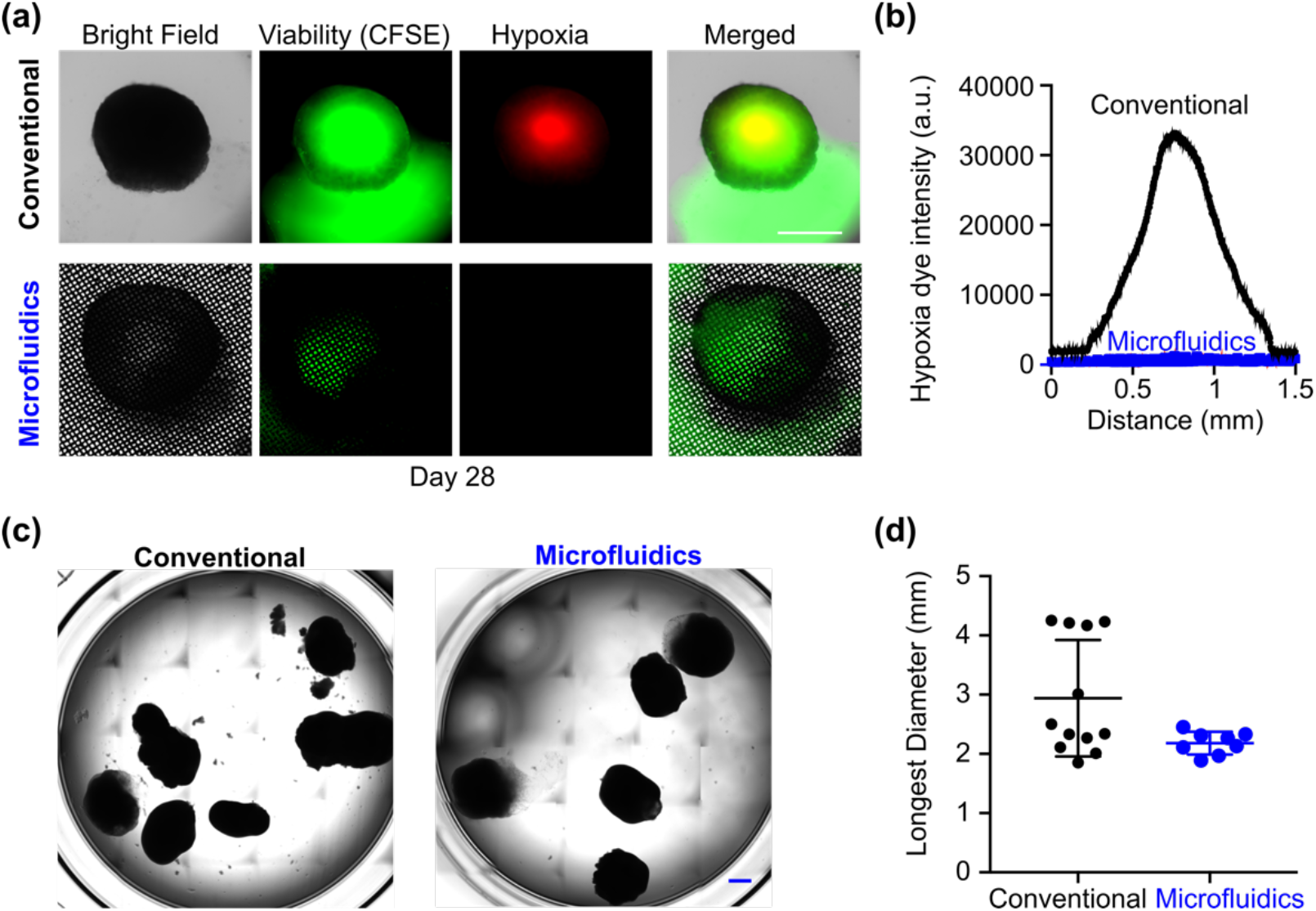
(a) Visualization of brain organoid hypoxia at 28 days in culture by conventional and microfluidic culture. (b) Quantification of hypoxia dye distribution across the brain organoids generated using conventional (top, in black) and microfluidic (bottom, in blue) methods. (c) Bright-field images of brain organoids fabricated using the conventional method and the microfluidic method. (d) The size distribution of brain organoids fabricated using the conventional method and the microfluidic method. Scale bar: 1 mm.

### Organoid maturation

We further interrogated the biological features of our brain organoids using cryo-sections and immunofluorescence staining. As shown in **Fig. 4a**, mature brain organoids demonstrated the formation of the ventricular zone (VZ) like areas indicated by paired box protein 6 (PAX6) staining of neural progenitor cells (NPC) and subventricular zone (SVZ) like areas indicated by microtubule-associated protein 2 (MAP2) staining. Furthermore, to confirm our brain organoids can respond to cannabinoid treatment, we also stained them for CB1 cannabinoid receptor (CB1) expression. As shown in **Fig. 4b**, many brain organoid cells express CB1, and it is enriched outside of VZ areas in post-mitotic neurons, similar to the pattern observed during rodent and human neonatal brain development^55–57^. Furthermore, we also observed intrinsic astrocyte development within our brain organoids indicated by staining with the astrocyte marker, glial fibrillary acidic protein (GFAP), which is also an essential component in the recapitulation of neonatal endocannabinoid signaling (**Fig. 4c**).^58^

**Figure 4.**
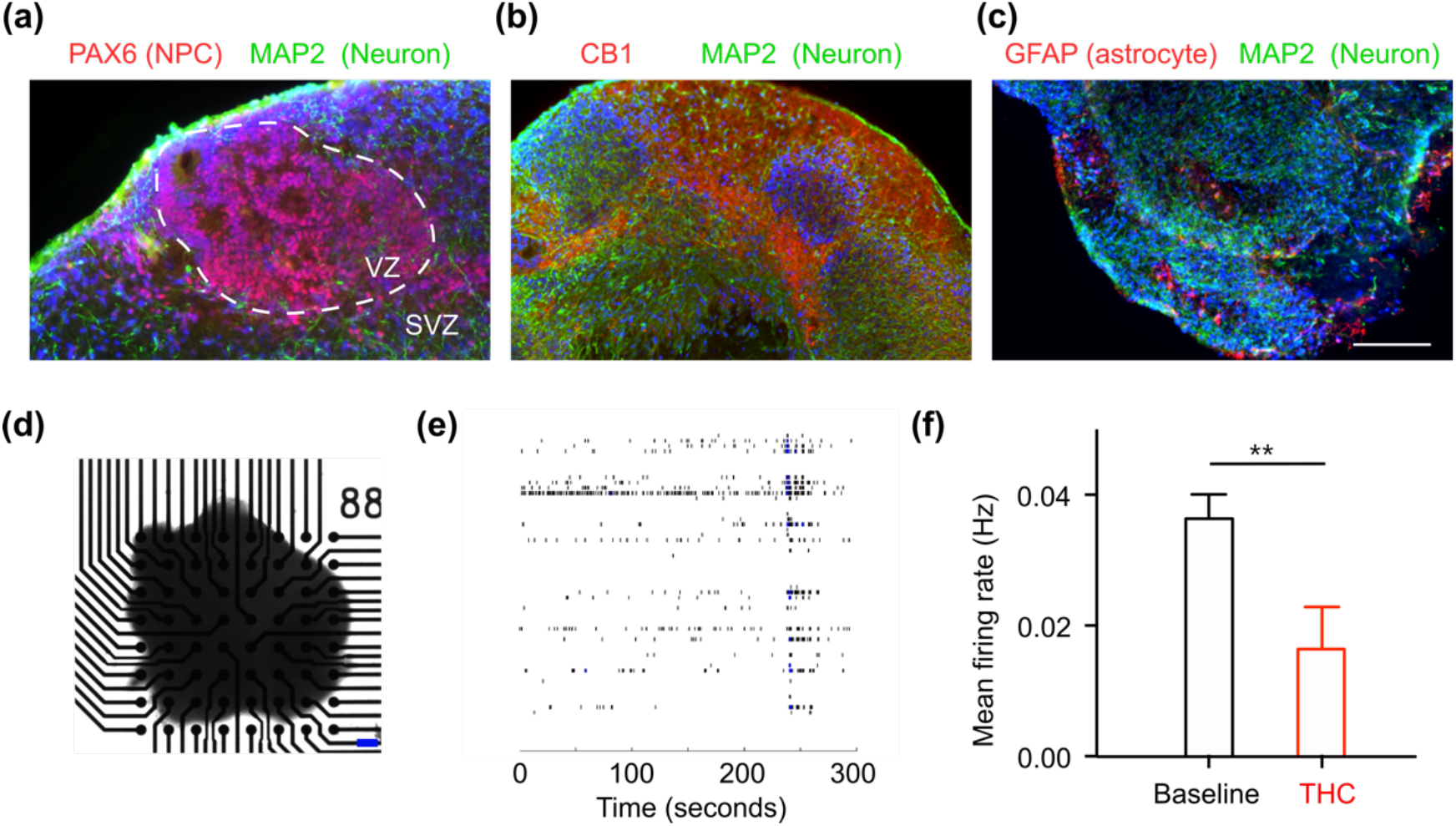
(a) Immunofluorescence staining of ventricular zone/subventricular zone formation in a brain organoid: PAX6 (neural progenitor cell (NPC), red), MAP2 (neuron, green), and DAPI (nucleus, blue). (b) Immunofluorescence staining of CB1 positive cells and their processes in a brain organoid: CB1 (red), MAP2 (neuron, green), and DAPI (nucleus, blue). (c) Immunofluorescence staining of astrocytes in brain organoid: GFAP (astrocyte, red), MAP2 (neuron, green), and DAPI (nucleus, blue). (d) Mature brain organoid attached to the MEA plate. (e) Example raster plot of brain organoid spontaneous activity. Each black tick mark indicates a neuronal firing on an electrode, and each row shows activity for one electrode. (f) Mean firing rate (Hz) of brain organoids (n=3) before and during THC treatment. Scale bar: 200 μm.

### Electric activity of mature organoids

We also characterized the electrical activity of our brain organoids and their responses to THC treatment. We measured their spontaneous firing and firing under delta-9-THC treatment using a microelectrode array (MEA) system. Briefly, mature brain organoids were allowed to attach to the MEA electrodes for 4 - 6 days before we recorded its spontaneous firing (**Fig. 4d**). Brain organoids were spontaneously active and exhibited burst activity (**Fig. 4e**). We then subjected the mature organoids to delta-9-THC treatment. As shown in **Fig. 4f**, the mean firing rate was significantly reduced with THC treatment (p < 0.01, n = 3), in concordance to that observed in mouse hippocampal slices^7^. This result confirms that our brain organoids can functionally react to transient delta-9-THC treatment.

### Modeling PCE in microfluidics

After characterizing our brain organoids with good activity, we then tested the effect of prolonged PCE on brain organoids. Our microfluidic device is good at performing medium changes and controlling THC perfusion of organoids. Briefly, after EB formation for 3 days, we started to treat the experimental group with 100 nM of delta-9-THC dissolved in DMSO, whereas the control group was dosed with vehicle only (0.1% DMSO). Brain organoids were treated for 27 days under prolonged THC dosing with the medium change from the bottom chamber of the microfluidic device every other day. At day 30 of THC treatment, brain organoids were analyzed by immunofluorescence microscopy as well as lysed for qRT-PCR. To analyze whether prolonged THC treatment affected neurodevelopment, we stained brain organoids for PAX6 and MAP2. Increased PAX6+ NPC number and VZ-like layer thickness were seen in the prolonged THC treatment group (**Fig. 5a**), this was likely due to the activation of CB1 receptors by delta-9-THC, which in turn stimulated NPC proliferation. This observation was also confirmed by the qRT-PCR results (**Fig. 5b**). Expression of the NPC marker, PAX6, was significantly increased in THC treated organoids, whereas the mature neuronal markers, CTIP2 and TUJ1 were decreased in THC-treated group (n = 5). This result indicated that prolonged THC treatment impacts human NPC proliferation as well as brain organoid structure. Furthermore, we also analyzed the impact of prolonged THC treatment on CB1 expression in brain organoids. As shown in **Fig. 5c**, CB1 expression was downregulated in the prolonged THC treatment group. This result was confirmed by the qRT-PCR results (n = 5) (**Fig. 5d**). This result is in concordance with previous human positron emission tomography (PET) scan results using a radioligand of CB1 in humans chronically consuming cannabis^59^. Lastly, as THC exposure was reported to reduce neurite outgrowth,^60^ we also tested if this phenomenon could be recapitulated using brain organoids *in vitro*. Mature brain organoids were allowed to adhere to a glass coverslip for 3 days in the presence or absence of 1 μM THC. We then stained outgrowths from neurons with cell tracker dye to visualize neurites. As shown in **Fig. 6**, neuron density, neurite length, and coverage areas were all decreased in the THC treated group. This indicated that THC treatment impairs neurite outgrowth in brain organoids.^61^

**Figure 5.**
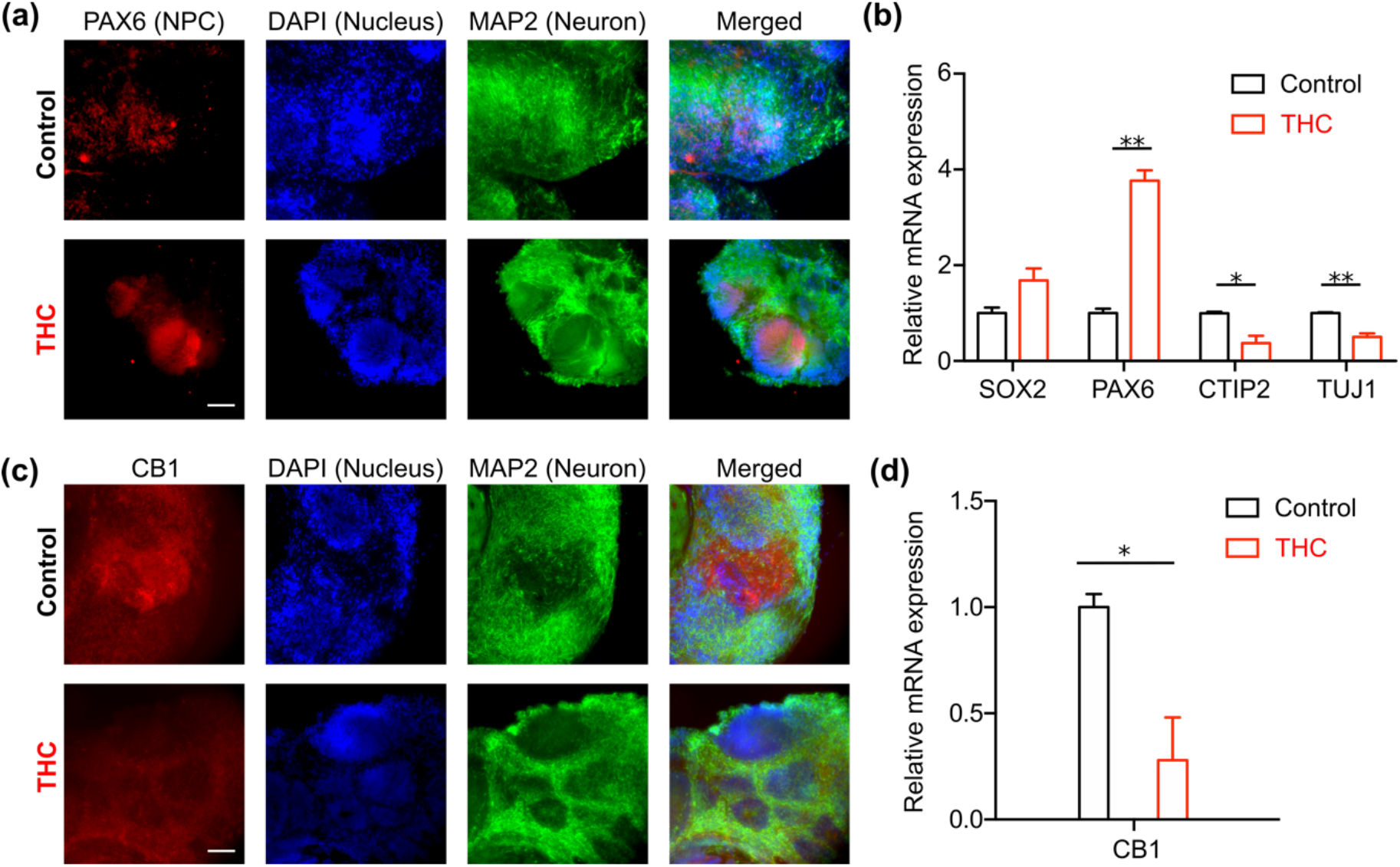
(a) Immunofluorescence staining characterizing the ventricular zone/subventricular zone distribution in brain organoids with and without THC treatment: PAX6 (neural progenitor cell (NPC), red), MAP2 (neuron, green), and DAPI (nucleus, blue). (b) relative quantitative PCR results analyzing neural progenitor and neuronal marker expression in brain organoids with and without THC treatment, n=5. (c) Immunofluorescence staining of CB1 protein in brain organoids with and without THC treatment, CB1 (red), MAP2 (neuron, green), and DAPI (nucleus, blue). (d) Relative quantitative PCR results to quantify CB1 expression in brain organoids with and without THC treatment, n=5. Scale bar: 100 μm.

**Figure 6.**
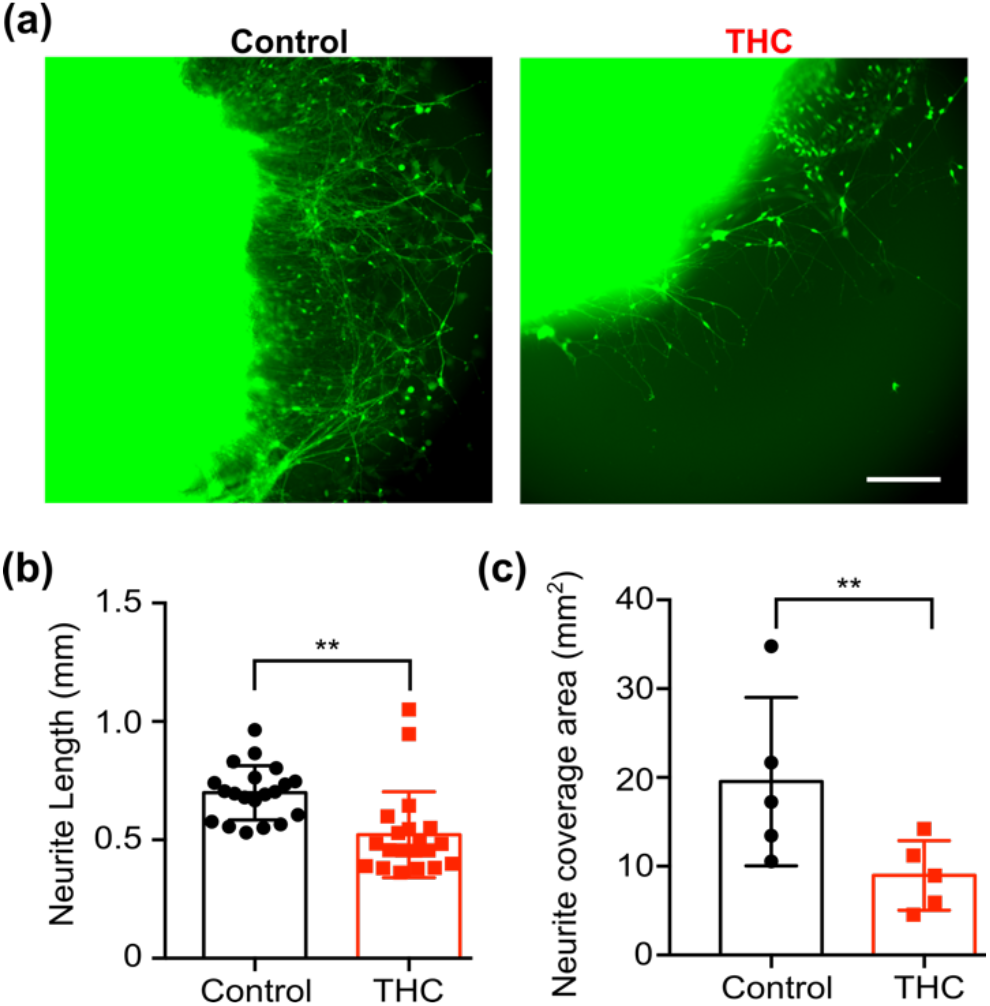
(a) Neurite outgrowth during control and THC treated conditions. (b) Quantification of neurite length from control and THC treated brain organoids. (c) Quantification of neurite coverage area per brain organoid after 3 days of adherent culture, n=5. Scale bar: 500μm.

## CONCLUSIONS

Here, we demonstrated a microfluidic device for a one-stop brain organoid assembly. This device minimized brain organoid size heterogeneity by preventing organoids from merging during the culture process and prevented losses associated with transferring the cultures. This device also integrated an air-liquid interface culture method to reduce hypoxic core formation in brain organoids. Utilizing this methodology, we analyzed the effect of prenatal cannabis exposure on brain development. We demonstrated that prolonged THC exposure altered the neonatal brain VZ/SVZ ratio, CB1 expression, neurite outgrowth, and spontaneous neuronal activity. The advantages of convenient one-stop fabrication and organoid uniformity control by this device allowed us to study the effects of THC on prenatal brain development while minimizing interference from internal brain organoid heterogeneity. The result of these preliminary studies shows that it is feasible to measure the impact of prenatal THC exposure on early neural developmental using cerebral organoids. Future studies are underway to investigate how THC can alter neural circuitry formation and functions using brain region-specific organoid models. Moreover, cannabis contains over 400 chemical compounds and at least 61 types of cannabinoids^62^. Other well-known cannabinoids such as cannabidiol (CBD) may also have an impact on early neural development. To comprehensively understand the effect of PCE, more studies are needed to fully understand the effect of the potential interaction of multiple cannabinoids and other chemicals (e.g., terpenes) present in cannabis in brain development. To conclude, we developed a simple-to-use, one-stop microfluidic device for brain organoid fabrication, modeling early human neuronal development. We used this device to study the effect of PCE on human brain development. In the future, we envision that this device can be adopted in a broad area of studies modeling early human brain development with the scalable capacity to accommodate the needs of pharmacology research.

## Acknowledgments

This project was supported by the departmental start-up fund of Indiana University Bloomington.

